# RNA degradomics and proteomics reveal the mechanism of dsProsβ1-mediated proteasome targeting in the cabbage stem flea beetle

**DOI:** 10.1101/2025.03.23.642439

**Authors:** Doga Cedden, Gözde Güney, Michael Rostás, Stefan Scholten

## Abstract

**Background:** The cabbage stem flea beetle (CSFB, *Psylliodes chrysocephala*) is a major threat to oilseed rape crops. Management of CSFB has become increasingly challenging due to the European Union’s ban on neonicotinoids and the emergence of pyrethroid-resistant populations. Recently, RNA interference (RNAi) has shown potential as an environmentally friendly alternative for the management of CSFB, and proteasome subunits have been identified as very effective RNAi targets. However, the mechanism of action of proteasome-targeting RNAi strategies remains to be fully characterized at the molecular level in CSFB and other pests. Here, we used CSFB to investigate the mechanism of action of dsProsβ1, which is a double-stranded RNA targeting a proteasome subunit.

**Results:** RNA degradome sequencing identified siRNA-mediated cleavage events in the target transcript, with cleavage events occurring at higher rates between uracil-guanine and adenine-adenine pairs. RISC-bound small RNA sequencing confirmed the presence of mature siRNAs guiding these cleavage events while revealing discrepancies between siRNA abundance and cleavage patterns. Proteomics analysis identified changes in protein levels caused by proteasome inhibition, including an increase in mitochondria- and cytoskeleton-related proteins and a decrease in central dogma-associated proteins.

**Conclusion:** This study demonstrates that combining RNA degradomics, RISC-bound sRNA-seq, and proteomics is an insightful approach to investigating the mechanism of RNAi-based pest control at the molecular level. The insights gained from these methods can be used to enhance proteasome-targeting RNAi strategies against insect pests.

**Graphical Abstract:** 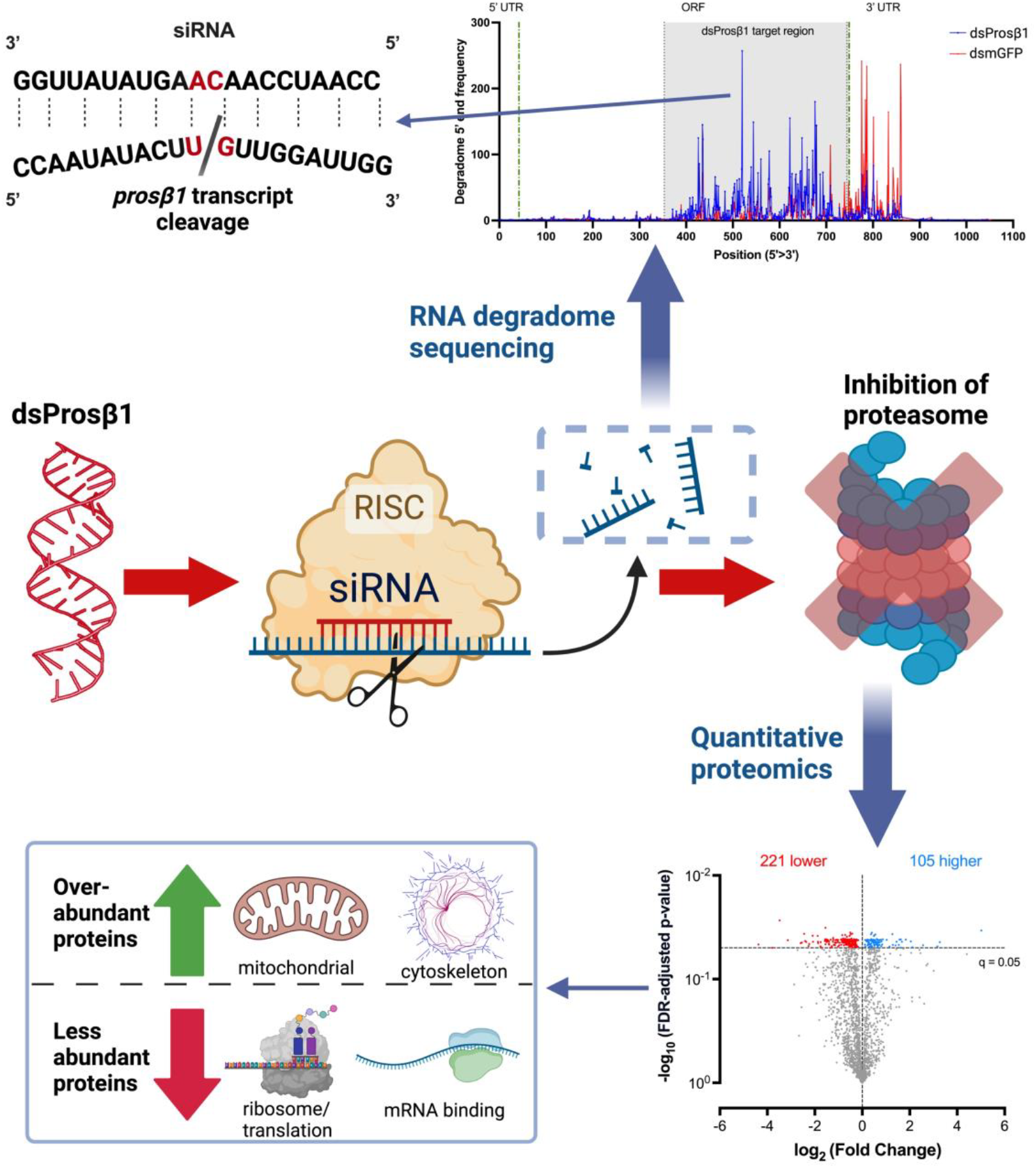

RNA degradomics revealed dsProsβ1-derived siRNA-mediated mRNA cleavage events, mainly at uracil-guanine and adenine-adenine pairs. Proteasome inhibition via dsProsβ1 increased mitochondrial and cytoskeletal proteins while reducing translation-related and mRNA-binding proteins.

## 1 Introduction

The damage inflicted by insect pests accounts for ∼20% of yield losses worldwide.^1^ Although chemical insecticides have allowed the control of pests, the emergence of insecticide-resistant populations and societal shift toward more environmentally friendly policies necessitates the development of alternative strategies.^2–4^ RNA interference has been recognized as a promising alternative to chemical insecticides in managing pest populations due to its environmentally friendly characteristics.^5,6^ RNAi-based pest management depends on the knockdown of essential target genes to cause insecticidal effects.^6–8^ Previous studies have identified very effective target genes, including those involved in the secretory pathway and proteasome-mediated protein degradation.^9–11^ Recently, a sprayable dsRNA that targets *proteasome subunit beta type-5* has been registered for the management of the Colorado potato beetle, *Leptinotarsa decemlineata*, highlighting the importance of proteasome in RNAi-based pest management.^12^ It is expected that the knockdown of proteasome subunits via dsRNA delivery compromises the protein degradation process and, consequently, leads to an unviable proteome and severe mortality.^13,14^ However, the mechanism of proteasome targeting insecticidal dsRNA remains to be understood at the molecular level.

RNAi is triggered by double-stranded RNA (dsRNA) produced by viruses and transposons in nature and by artificially produced dsRNA in the context of pest control.^15–17^ When the endonuclease Dicer-2 recognizes intracellular dsRNA, it cleaves dsRNA into ∼19 base pair RNA duplexes with two-nucleotide 3′ overhangs, known as small interfering RNA (siRNA).^18– 20^ One of the strands of siRNA duplex is selected as the guide strand and guides the RNA-induced silencing complex (RISC) to cleave complementary RNA, resulting in sequence-specific silencing.^21–23^

The cabbage stem flea beetle (CSFB) is a notorious pest of winter oilseed rape, especially in northern Europe.^24,25^ The control of CSFB has become very challenging due to the ban on neonicotinoid treatments in Europe and the emergence of pyrethroid-resistant pest populations.^26,27^ Previous research demonstrated the potential of RNAi in the control of CSFB adults^28^ and subsequently, proteasome subunits were identified as very effective RNAi targets.^9^ The feeding of dsRNA targeting selected proteasome subunits, including Pc-*prosβ1*, led to 100% mortality within 2 weeks and up to 75% feeding inhibition within 5 days.^9^

RNA degradome sequencing, also known as parallel analysis of RNA ends (PARE) or global mapping of uncapped and cleaved transcripts (GMUCT), has been recognized as a valuable tool for identifying small RNA-mediated cleavages in plants.^29–31^ Identification of the cleavage events is state of the art in demonstrating transcript regulation by various types of small RNA.^29^ However, RNA degradome sequencing has not been utilized to characterize the cleavage events caused by insecticidal dsRNA, which may give additional insights into its mode of action and inspire strategies for optimizing RNAi-based strategies. For instance, RNA degradome sequencing might help us identify local differences within the delivered dsRNA in causing target mRNA cleavages. If specific dsRNA sequence motifs have a higher potential to cause target mRNA cleavages, this knowledge can guide designing dsRNA with higher insecticidal efficacy. Indeed, it has been shown that consideration of siRNA sequence features improves insecticidal efficacy of essential gene targeting dsRNA in insect pests and the dsRIP web platform was developed for this purpose.^32^

Here, we used CSFB as a pest model^33^ to study the mechanism of action of a proteasome targeting insecticidal dsRNA (dsProsβ1). We performed RNA degradomics to identify the siRNA-mediated cleavage events and sequenced RISC-bound small RNA to identify mature siRNAs processed from orally delivered dsProsβ1. Furthermore, we conducted proteomics analysis to study the changes in the protein levels caused by the inhibition of the proteasome by dsProsβ1.

## 2 METHODS

### 2.1 Insect culture

The laboratory colony of CSFB (*Psylliodes chrysocephala*) was kept in rearing chambers set to 21° ± 1°C and 65 ± 10% relative humidity under a 16:8 light: dark regime. The beetles were reared on oilseed rape plants (BBCH-scale: 30–35). Freshly emerged adults were collected daily for bioassays or returned to the rearing chambers to maintain the colony. To preserve genetic diversity, CSFB larvae collected from a pesticide-free experimental oilseed rape field in Göttingen, Germany (coordinates: 51.564065, 9.948447) were annually introduced into the laboratory colony.

### 2.2 Bioinformatic characterization

The sequence of Pc-*prosβ1* was retrieved from a CSFB adult de novo transcriptome assembly representing all three adult stages.^33^ The ORF and protein sequences were obtained using TransDecoder (v5.7, https://github.com/TransDecoder/TransDecoder/wiki). The 3D structure of the protein was predicted using AlphaFold 3 web server.^34,35^ The conserved domain analysis was conducted using NCBI Conserved Domain Search.^36^ The protein sequences orthologous and paralogous to Pc-Prosβ1 were retrieved from NCBI. The phylogenetic tree of protein sequences was constructed using the Neighbor-Joining method in MEGA 11 with 500 bootstrap replications.^37^

### 2.3 dsRNA synthesis and feeding

dsProsβ1 is a double-stranded RNA 397 bp in length and is complementary to the ORF of the Pc-*prosβ1* gene (for further details, see Tab. S1 and ^9^). A control dsRNA, dsmGFP, was synthesized from a DNA template (IDT, Germany)^9^. Both dsRNAs avoided off-targets by skipping regions with ≥21 bp matches to other transcripts in CSFB adults. Bowtie v1.1^38^ was used to identify potential off-targets as described previously by Cedden et al. (2024)^10^. Primer3^39^ with default settings was used to design primers to amplify the target region, and T7 promoter sequences (‘GAATTGTAATACGACTCACTATAGG’) were added to the 5′ ends of the primers (IDT, Germany). RNA was extracted from mixed-age CSFB adults and reverse transcribed into cDNA using Quick-RNA Tissue Kit (Zymo Research) and LunaScript® RT SuperMix Kit (NEB), respectively. To obtain the DNA template for in vitro dsRNA transcription, two separate PCR reactions were performed with primer pairs containing T7 promoter in either the forward or the reverse primer and the CSFB cDNA using Q5® High-Fidelity Master Mix (NEB). Single PCR bands were purified for *in vitro* transcription using the MEGAscript™ T7 Kit. Forward and reverse RNA strands were synthesized in separate 20 µL reactions and then equimolecularly combined following lithium chloride precipitation-based RNA purification. The dsRNA was denatured at 94 °C for 5 min and reannealed at 25 °C for 30 min, and the length was verified by running 1% agarose gel.

dsRNAs were diluted to 500 ng/μL in nuclease-free water containing Triton-X (200 ppm final concentration, Thermo Fisher Scientific) and 1 µL of dsRNA solution was homogenously pipetted on a 30 mm^2^ leaf disk punctured from the first true leaves of oilseed rape plants. Individual newly emerged CSFB females were placed into Petri dishes (60 × 15 mm with vents) together with the dsRNA-containing leaf disk, which was placed on 1% agarose gel to preserve freshness. Untreated 400 mm^2^ fresh leaf disks were placed into the Petri dishes on the second day of the bioassay (when the dsRNA-treated leaf disk was consumed) and subsequently every third day.

### 2.4 RNA degradome sequencing

dsRNA-fed CSFB were sampled 3 days post dsRNA treatment for RNA degradome sequencing as previously established.^40^ Total RNA was isolated from 20 adults fed with dsProsβ1 or dsmGFP using the Quick-RNA Tissue Kit (Zymo Research). RNA quality was evaluated with an Agilent 2100 Bioanalyzer, and 100 µg of RNA was used for degradome library preparation. Polyadenylated RNA was purified with Dynabeads mRNA Purification Kit and Oligo d(T)25 Magnetic Beads. A 5’ RNA adaptor (100 µM) was ligated using T4 RNA ligase, and adaptor-ligated RNA was repurified. First-strand cDNA synthesis was performed with ProtoScript II Reverse Transcriptase (NEB). The second-strand was synthesized using MyFi Mix and 5’/3’ adaptor primers (Tab. S2) with 15 PCR cycles. The PCR product was purified, digested with EcoP15I (NEB), and ligated to dsDNA adaptors (Tab. S2). The resulting 79 bp DNA was purified through PAGE and concentrated via ethanol precipitation. Thirteen PCR cycles with MyFi Mix and library-specific primers added indexes to the libraries (Tab. S2). Libraries were pooled, PAGE-purified, and sequenced on the DNBSEQ-G400 platform (PE50+5+10 mode) by BGI Tech Solutions (Hong Kong).

The raw reads were trimmed and quality-filtered (q > 30) using cutadapt (v4.4, https://journal.embnet.org/). dsProsβ1 sequence (Tab. S1) was converted into all possible 21-mer siRNA sequences in either direction. CleaveLand4 (v4.5) with the default parameters was used to analyze the RNA degradomics data. The degradome frequency outputs from dsProsβ1- and dsmGFP-fed samples were plotted using Graphpad v10.

### 2.5 RISC-bound small RNA sequencing

dsRNA-fed CSFB were sampled 3 days post dsRNA treatment for RISC-bound small RNA sequencing as previously established.^32^ Small RNA was isolated from four CSFB whole bodies using the TraPR Small RNA Isolation Kit (Lexogen GmbH, Vienna, Austria).^41^ Samples were homogenized in 300 µL lysis buffer with RNAse-free pestles, vortexed for 20 seconds, and centrifuged (10,000 g, 5 min at 4°C). Cleared lysates were loaded onto TraPR columns mixed with resin, and RISC was eluted through centrifugation (1,000 g, 15 sec, repeated thrice). The final RISC elution (750 µL) was subjected to RNA precipitation using phenol (pH 4.3) / chloroform / isoamylalcohol pre-mix (ROTI®Aqua-P/C/I for RNA extraction, Carl Roth GmbH, Germany) and 3M sodium acetate and and carrier substances (TraPR kit) to facilitate the precipitation. The RNA pellets were washed with 80% ethanol and eluted in 10 µL RNA elution buffer. The RISC-bound small RNA sequencing experiment was repeated twice.

Small RNA libraries were prepared using the NEBNext® Multiplex Small RNA Library Prep Kit (New England Biolabs GmbH, Germany). Six microliters of small RNA (∼200 ng) was used as input. Adaptors for Illumina were not diluted due to the high RNA input, and libraries were amplified by 11 PCR cycles. The amplified DNA was purified with the Monarch PCR & DNA Cleanup Kit and size-selected (135–150 bp) through 6% polyacrylamide gel. Libraries were quality-checked with an Agilent 2100 Bioanalyzer, pooled, and sequenced on the DNBSEQ-G400 platform by BGI Tech Solutions (Hong Kong).

Raw reads were trimmed and quality-filtered (q > 30) using TrimGalore (v0.6) with default parameters. Cleaned reads were mapped to the respective dsRNA sequences using Bowtie 1 (v1.3) with the parameters “-all” and “n=0” to allow all valid forward and reverse mappings with no mismatches. Mapped 21 nt read counts were extracted using SAMtools (v1.2, https://www.htslib.org) and percent normalized, where 100% corresponds to total read count mapping onto the dsRNA. Degradome frequency at the location corresponding to the 11^th^ nucleotide of the complementary siRNA was considered during the integration of the two datasets. Pearson correlation was calculated using GraphPad v10.

### 2.6 Proteomics

Whole bodies of four CSFB adults receiving dsRNA treatment were sampled on 5 days post treatment, frozen in liquid nitrogen, and homogenized by grinding. Proteins were extracted with a urea denaturation buffer (6 M urea, 2 M thiourea, 10 mM HEPES) at a ratio of 1 mL per 100 mg sample. Supernatants were recovered by centrifugation (12,000 g, 5 min), and protein concentrations were measured using a Bicinchoninic Acid (BCA) assay. Proteins were precipitated using a chloroform/methanol method, digested using trypsin (Thermo Fisher Scientific) and RapiGest SF solution (Waters), treated with trifluoroacetic acid (1:10), and incubated at 37°C for 45 minutes before drying in a speed vac. Samples were purified using C18 stage tip protocol^42,43^ and analyzed at the Service Unit LC/MS Protein Analytics (Göttingen Center for Molecular Biosciences). Briefly, LC/MS analysis of the peptide samples was performed using an Ultimate 3000 system (Thermo Fisher Scientific) coupled to a Q Exactive HF mass spectrometer (Thermo Fisher Scientific). Peptides were separated by reverse phase liquid chromatography and on-line ionized by nano-electrospray (nESI) via the Nanospray Flex Ion Source (Thermo Fisher Scientific). Scans were recorded in a mass range of 300-1650 m/z (30,000 resolution). The XCalibur 4.0 software (Thermo Fisher Scientific) was used for LC-MS method programming and data acquisition. The proteomics experiments were repeated five times.

Label-free quantification (LFQ) values were obtained by mapping peptides to the CSFB adult proteome using MaxQuant (https://www.maxquant.org).^33^ Proteins with fewer than three peptide counts were discarded. Welch’s test was conducted and statistical significance was set at adjusted P < 0.05 (permutation-based false discovery rate), along with LogFold_2_ > 1 (increase in abundance) or LogFold_2_ < -1 (decrease in abundance) using Perseus (https://maxquant.net/perseus/).^44^ Proteins significantly changing in abundance were subjected to Gene Ontology (GO) enrichment analysis using the R package clusterProfiler (v3.17). The functional annotations of the proteins were retrieved from our CSFB annotation produced using Trinotate (v4.0, https://github.com/griffithlab/rnaseq_tutorial/wiki/Trinotate-Functional-Annotation).^33^

## 3 RESULTS

### 3.1 Bioinformatic characterization of Pc-*prosβ1*

Our previous RNAi screen identified Pc-*prosβ1* as a very effective RNAi target, causing not only 100% mortality within 2 weeks but also feeding inhibition up to 75% upon the oral delivery of 500 ng dsProsβ1.^9^ Here, we performed a bioinformatic characterization of *Pc-prosβ1*. Of note, the Pc-*prosβ1* gene was misannotated by the automated pipeline used in our previous screen-based study as *Pc-prosβ6*.^9^ The in-depth bioinformatic characterization revealed that this gene was a Beta-1 ortholog, rather than Beta-6 (Fig. 1B and Fig. S1). *Pc-prosβ1* contains an ORF spanning 702 nucleotides and encodes for 234 amino acids. The prediction of the 3D structure of Pc-Prosβ1 suggested a typical globular folding formed by 4 alpha sheets flanked by beta sheets (Fig. 1A). This structure is likely important for forming the internal Beta ring structure of the proteasome complex through interactions with other proteasome subunits. Conserved domain analysis suggested N-terminal nucleophile hydrolase (Ntn hydrolase) activity for Pc-Prosβ1, which is expected of a Beta 1 subunit. The Pc-Prosβ1 clustered with Prosβ1 orthologs of other coleopteran pests rather than other insect pest orders (Fig. 1B) and Prosβ1 was highly conserved among coleopteran pests at the protein sequence level (Fig. 1C).

**Figure 1.**
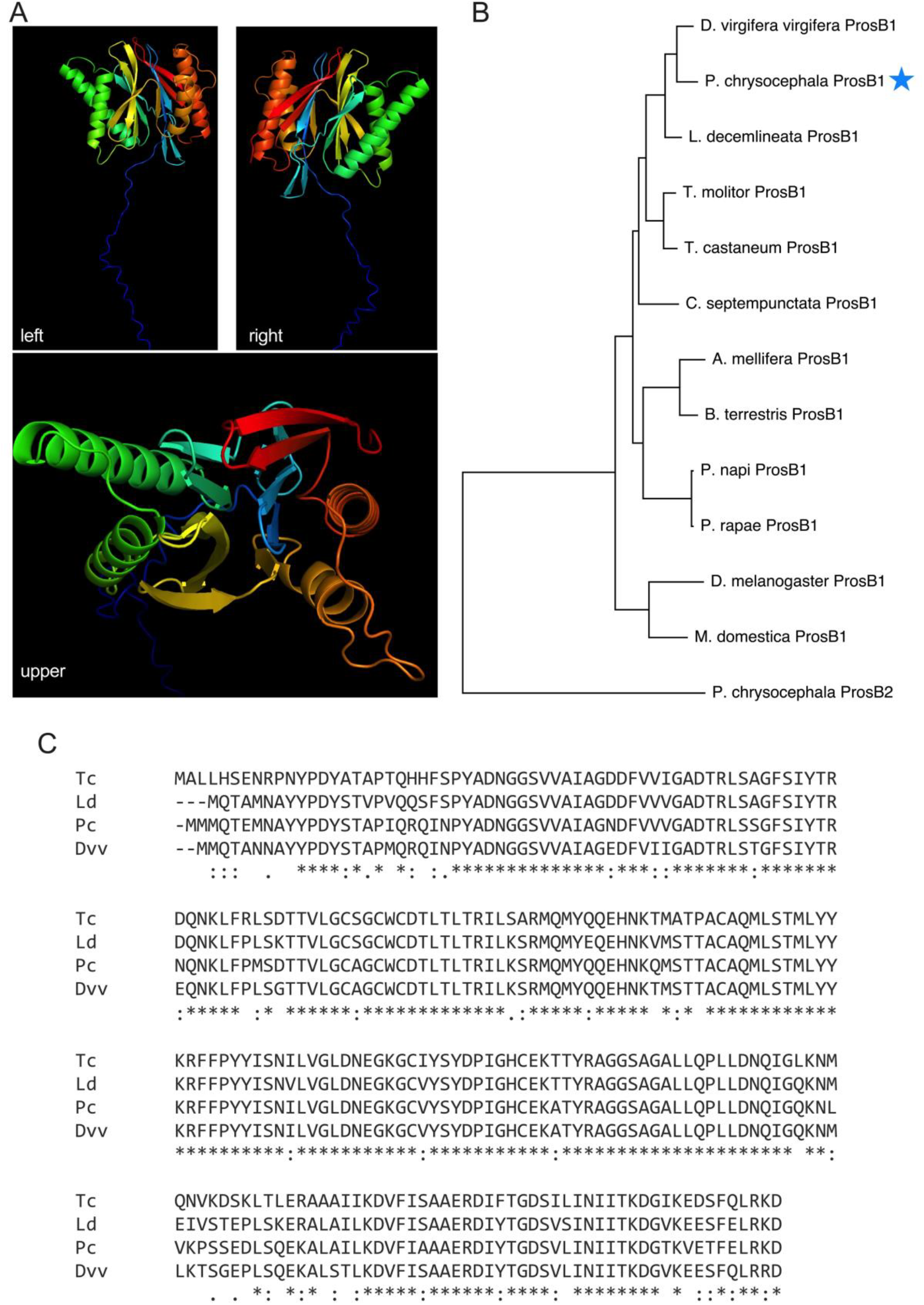
Bioinformatic characterization of *Pc-prosβ1* proteasome subunit. (A) The prediction of the 3D structure of Pc-Prosβ1 by AlphaFold 3 (https://alphafoldserver.com/) suggested a typical globular folding formed by 4 alpha sheets flanked by beta sheets. (B) Phylogenic tree of Prosβ1 orthologs from different insect species and CSFB paralog (Prosβ2) constructed using Neighbor-Joining method in MEGA 11 with 500 bootstrap replications. (C) Alignment of Prosβ1 orthologs from *Tribolium castaneum* (Tc), *Leptinotarsa decemlineata* (Ld), *Psylliodes chrysocephala* (Pc), and *Diabrotica virgifera virgifera (*Dvv) using Clustal Omega (https://www.ebi.ac.uk/jdispatcher/msa/clustalo, EMBL-EBI).

### 3.2 RNA degradome identifies cleavages caused by dsProsβ1

RNAi causes the cleavage of RNA transcripts complementary to dsRNA. Here, we performed RNA degradome sequencing in CSFB adults that were sampled 3 days after feeding on an oilseed rape disk containing 500 ng dsProsβ1 or dsmGFP. As expected, we identified various major cleavage events in the *Pc*-*prosβ1* transcript, specifically within the dsProsβ1 target region (positions 353 to 749 nt on the transcript), in the dsProsβ1-fed group (Fig. 2A). However, some cleavages in *Pc*-*prosβ1* transcript were also observed in the dsmGFP group, which is attributable to endogenous transcript processing. The cleavage sites in the dsmGFP group were more abundantly observed in the 3’ UTR of the *Pc-prosβ1*, suggesting endogenous regulation of *Pc-prosβ1* transcript mainly includes 3’ UTR cleavages, which is commonly observed in the form of endonucleolytic cleavage for polyadenylation and miRNA-mediated post-transcriptional regulation in animals.^45,46^ dsmGFP-fed CSFB did not contain any small RNA reads fully complementary to the 3’ UTR region of *Pc-prosβ1*; however, we found one miRNA potentially targeting the 3’ UTR region of *Pc-prosβ1*, which may be responsible for these cleavage events in control beetles (see Tab. S3, data from ^40^).

**Figure 2.**
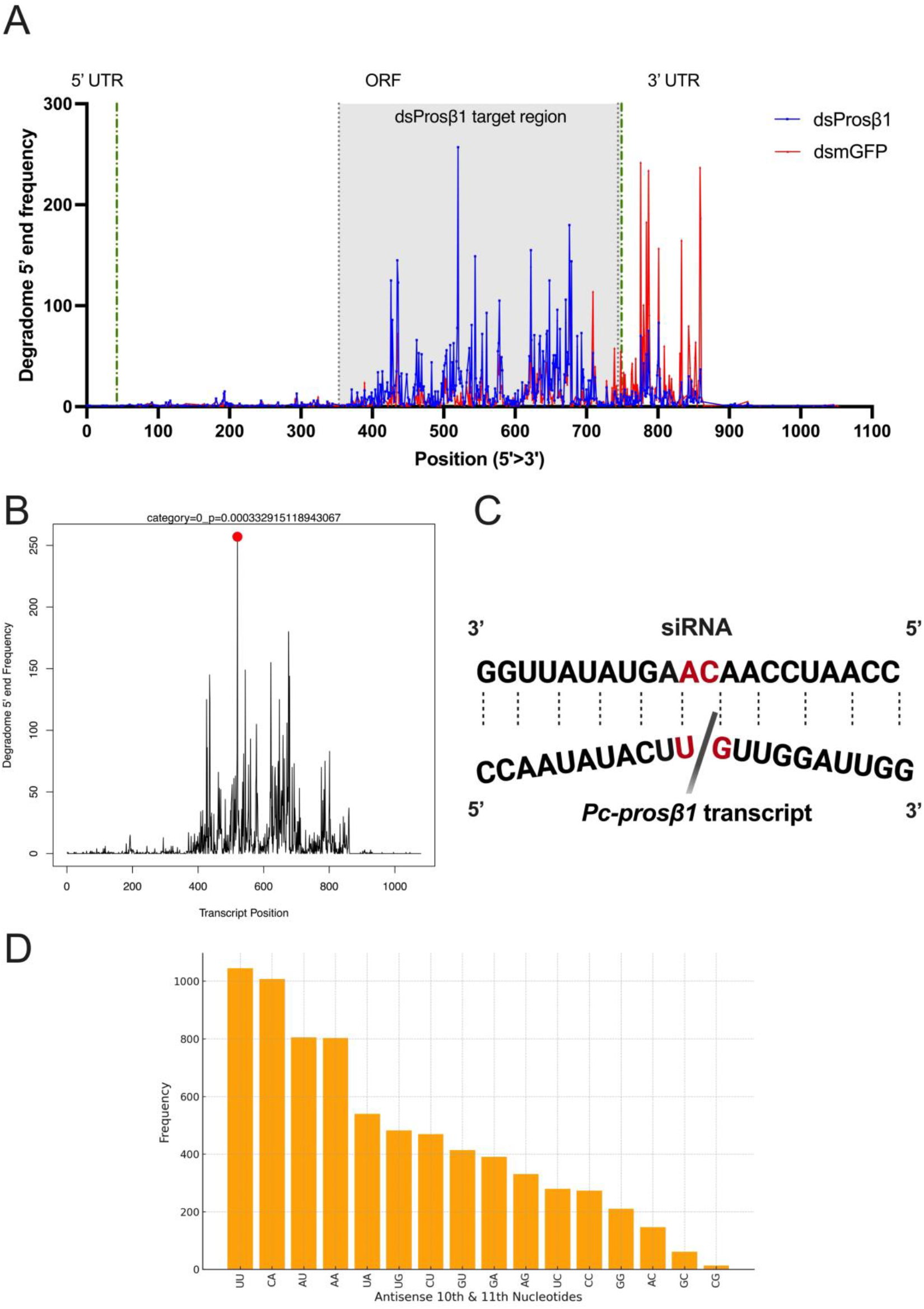
RNA degradomics identifies the cleavage sites on the *Pc-prosβ1* transcript. Newly emerging cabbage stem flea beetle (CSFB) adults were fed on oilseed rape leaf disk containing 500 ng of dsProsβ1 which is complementary to positions 353 to 749 bp in Pc-*prosβ1* transcript or dsmGFP (control dsRNA) and sampled 3 days after the consumption of the leaf disk (for this analysis RNA from 20 beetles was pooled into a single replicate). (A) Degradome 5’ end frequency (i.e., cleavage site frequency) plot of Pc-*prosβ1* transcript. Blue values indicate the cleavages identified in dsProsβ1-fed CSFB and red values indicate those identified in dsmGFP fed CSFB. (B) CleaveLand4 (v4.5) output with the most abundantly detected cleavage event (red dot) in dsProsβ1 fed CSFB. Category “0”, events with highest evidence are shown. (C) Representation of the most abundantly detected cleavage event in dsProsβ1 fed CSFB. The scissile phosphodiester bond (11^th^ and 12^th^ nucleotides on target site, corresponding to the 10^th^ and 11^th^ nucleotides on the guiding siRNA strand). (D) Frequencies of 10^th^ and 11^th^ nucleotide pairs in the antisense siRNA corresponding to the cleavage events.

The major cleavage event in the *Pc-prosβ1* transcript in the dsProsβ1 group was not observed in the dsmGFP group, indicating it was a specific peak in the former group (Fig. 2A). This peak corresponded to 520 nt position on the transcript and 167 bp position on the dsProsβ1, which has a total length of 397 bp. The scissile phosphodiester bond was between a uracil and a guanine and was categorized as “0”, which indicates the peak with the highest cleavage evidence in a transcript by CleaveLand4 (Fig. 2B-C). This exact cleavage was observed 257 times in the degradome library and the minimum free energy (MFE) score was -32.6 (Fig. 1D). There were also other major cleavage sites in the dsProsβ1 group that corresponded to the dsProsβ1 target region, especially around 600 to 700 nt on the transcript (Fig. 2A). Interestingly, the sites near to the start and end of the dsProsβ1 target region had lower amount of cleavage events, suggesting the ends of dsRNA were not very effective at causing cleavages. Plotting of frequencies of 10^th^ to 11^th^ nucleotide pairs in the antisense siRNA corresponding to the cleavage events showed that uracil-uracil (14.3% compared to 6.25% if all pairs had equal frequencies), cytosine-adenine (13.8%), and adenine-uracil (11%) pairs had the highest frequencies, which correspond to adenine-adenine, uracil-guanine, and adenine-uracil pairs in the mRNA, respectively (Fig. 2D). We also noted that guanine-cytosine or cytosine-guanine as 10^th^ to 11^th^ nucleotide pairs in the antisense siRNA were underrepresented in the frequency analysis (Fig. 2D).

### 3.3 RISC-bound sRNA-seq identifies mature siRNAs processed from dsProsβ1

dsRNA is first processed into siRNA, which is loaded onto RISC to guide mRNA cleavages. To further characterize the mode of action of dsProsβ1, we isolated and sequenced RISC-bound sRNA to investigate the mature siRNA processed from dsProsβ1. There was a major peak at 21 nt length for siRNA mapping to ds*Prosβ1* (Fig. 3A). The nucleotide frequency plot for highly abundant antisense strands showed slight biases in favor of adenine at the 1^st^ postion, uracil at the 3^rd^ position and guanine at the 18^th^ and 19^th^ nt positions compared to lowly abundant antisense strands (Fig. 3B). We did not detect siRNA outside the dsProsβ1 target region (Fig. 3C), highlighting the specificity of our RISC-bound sRNA sequencing approach. Two major antisense siRNA peaks were observed between 400 and 500 nt on the *prosβ1* transcript (Fig. 3C).

**Figure 3.**
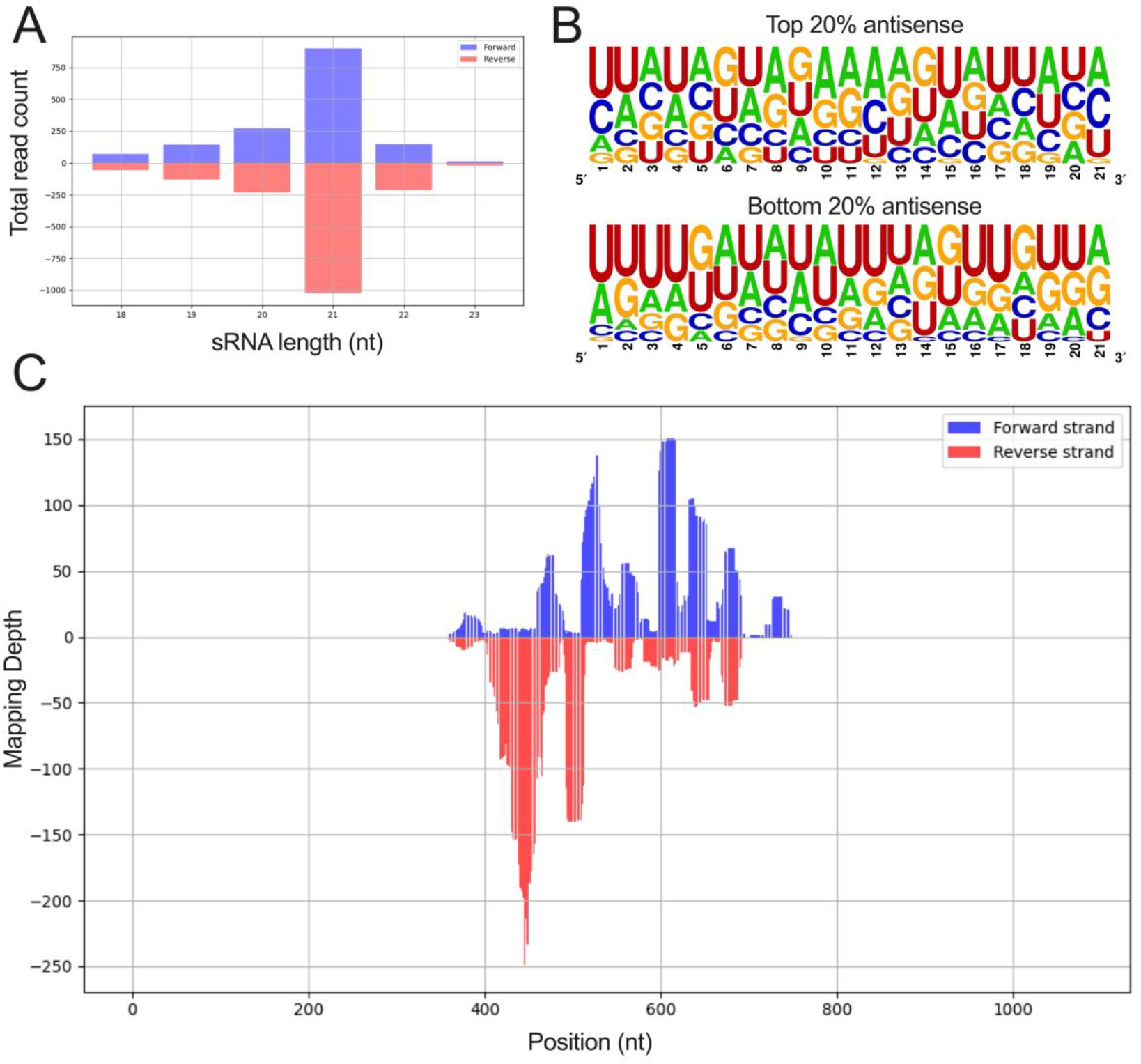
RISC-bound small RNA-seq identifies Pc-*prosβ1* targeting small interfering RNAs. Four newly emerging cabbage stem flea beetle (CSFB) adults were fed with oilseed rape leaf disk containing 500 ng of dsProsβ1 or dsmGFP (control dsRNA) and sampled 3 days after the consumption of the leaf disk (n = 2 per treatment). (A) Length distribution of dsProsβ1-mapping RISC-bound siRNA. (B) Plots showing the frequency of each nucleotide at each position in 21-nt long dsProsβ1-derived RISC-bound antisense siRNA. The top plot shows the top 20% antisense siRNA, while the bottom plot shows the bottom 20% antisense siRNA in terms of abundance (n = 43). (C) Mapping depth of 21-nt RISC-bound siRNA on the *Pc*-*prosβ1* mRNA sequence.

### 3.4 Integration of RNA degradome and RISC-bound sRNA-seq

Next, we extracted the 21-nt antisense RISC-bound siRNA generated from dsProsβ1 and matched them with the cleavage events identified through RNA degradome sequencing (Fig. 4). This analysis showed both convergences and discrepancies between the two data sets. For instance, the siRNA corresponding to the major cleavage event (at position 520) did not have a high amount of read counts in the RISC-bound siRNA data (Fig. 4A). Nonetheless, especially the first cleavage peak (position ∼440 on the transcript) and to some degree the last cleavage peak (position ∼760 on the transcript) were well covered by corresponding highly abundant RISC-bound siRNA (Fig. 4A and Tab. 1). Pearson correlation analysis between the degradome frequency and antisense siRNA abundance coinciding with the cleavage event showed a positive and significant correlation (P = 0.018), albeit the correlation coefficient was low (r = 0.224, Fig. 4A). As expected, Pearson correlation analysis between the degradome frequency and 21-nt sense siRNA abundance corresponding to the cleavage event was not significant (P = 0.94, r = -0.007). Overall, the integration of these data suggests that although there is the expected correlation between antisense siRNA abundance and cleavage events, there are likely other factors that cause discrepancies between the two data sets.

**Table 1.**
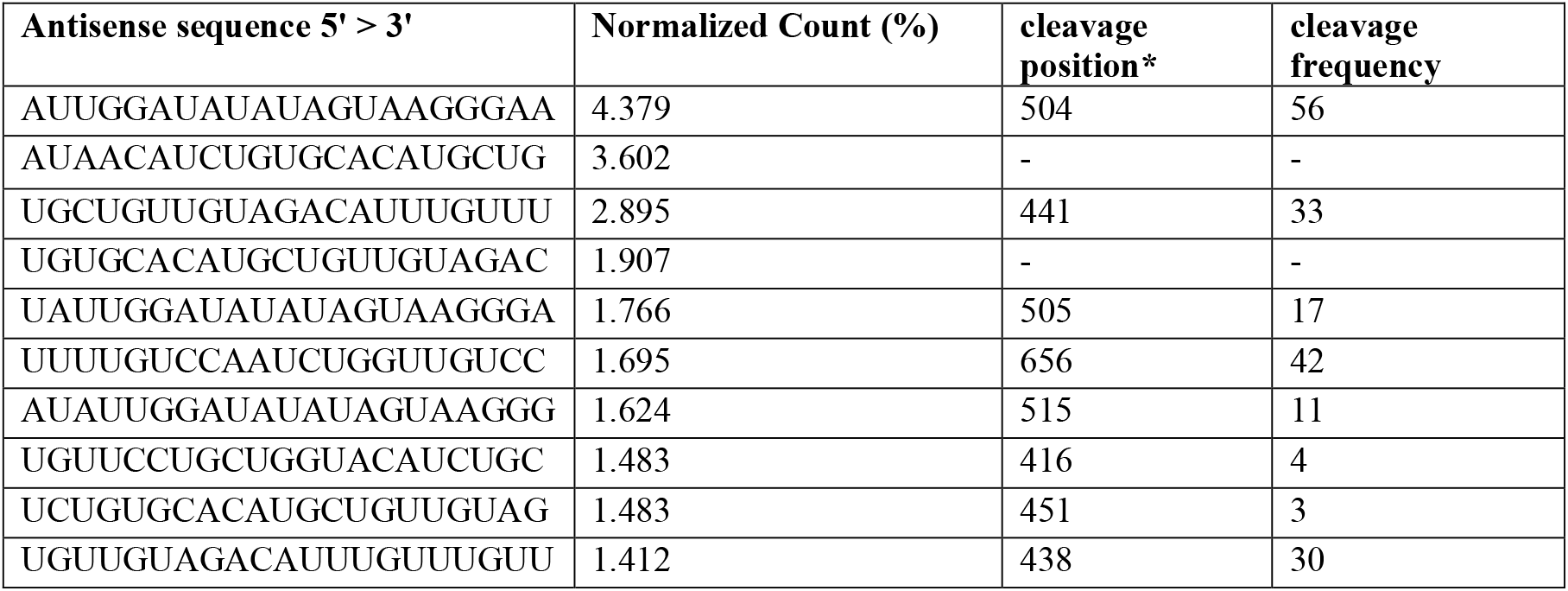
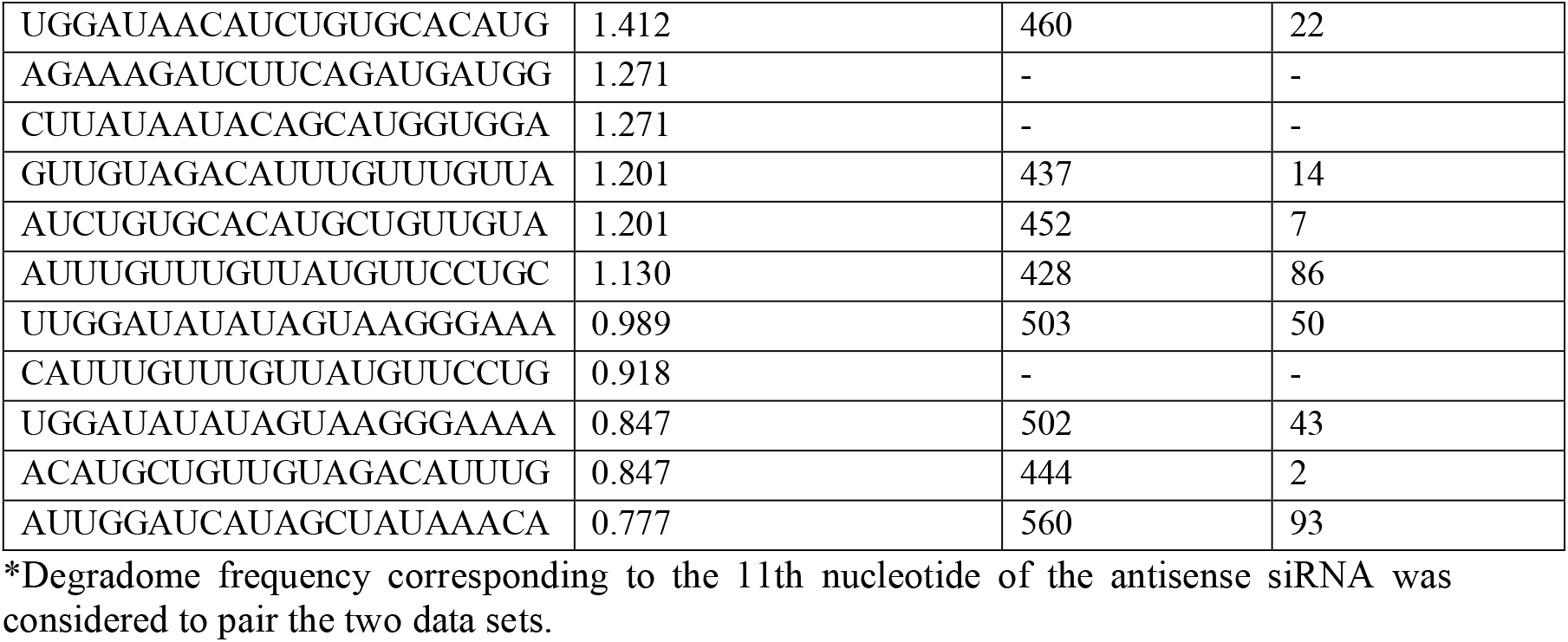
Top 20 most abundant 21-nt antisense RISC-bound siRNA reads mapping onto the delivered dsProsβ1 and respective cleavages in *Pc-prosβ1* mRNA.

**Figure 4.**
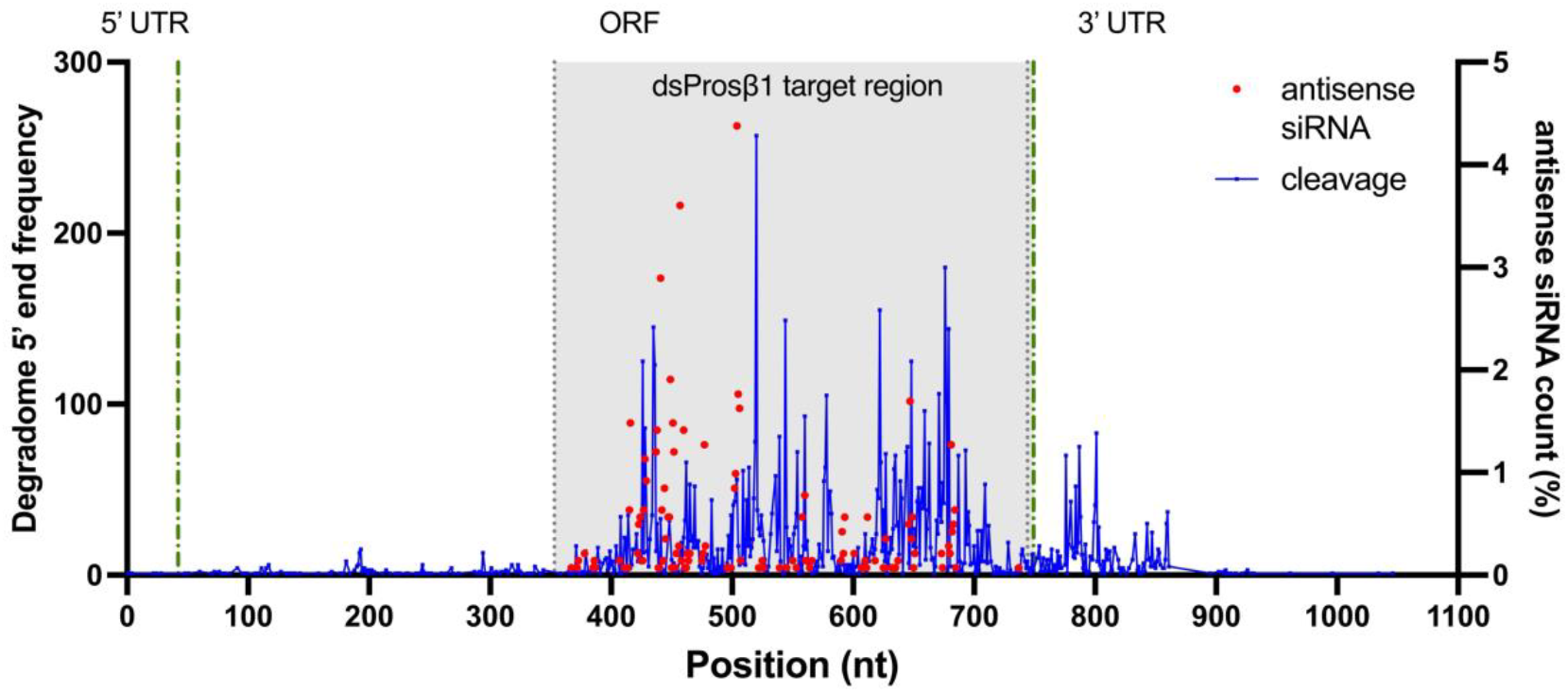
Combining RISC-bound siRNA abundance and RNA degradomics. Data presented in Fig. 1 and Fig. 2 were combined to plot the degradomics frequency per position on transcript together with corresponding RISC-bound antisense siRNA count. Pearson correlation coefficient was calculated between the degradomics frequency and RISC-bound antisense siRNA count (n = 86 pairs). Degradome frequency corresponding to the 11^th^ nucleotide of the antisense siRNA was considered to integrate the two data sets.

### 3.5 Proteomics reveals changes in protein levels caused by dsProsβ1

dsProsβ1 targets the proteasome, which is responsible for degrading unnecessary and misfolded proteins to recycle amino acids and prevent cellular stress and protein aggregation. Here, we compared the proteomes of dsProsβ1 and dsmGFP-fed CSFB to identify the changes in the protein levels caused by dsProsβ1. In total, we identified 326 proteins (out of 1980 proteins in total) showing differential abundance in the dsProsβ1 group (Fig. 5A). Of note, different isoforms of the same gene were counted as a single protein. In the dsProsβ1 group, the majority (221) of the differentially abundant proteins had lower abundance, while only around one-third (105) of the differentially abundant proteins had higher abundance. We also noted a skew towards lower abundance in the dsProsβ1group among proteins below the significance threshold, which further supports the idea that lowering, rather than increasing, the abundance of proteins is a more prominent effect of dsProsβ1.

**Figure 5.**
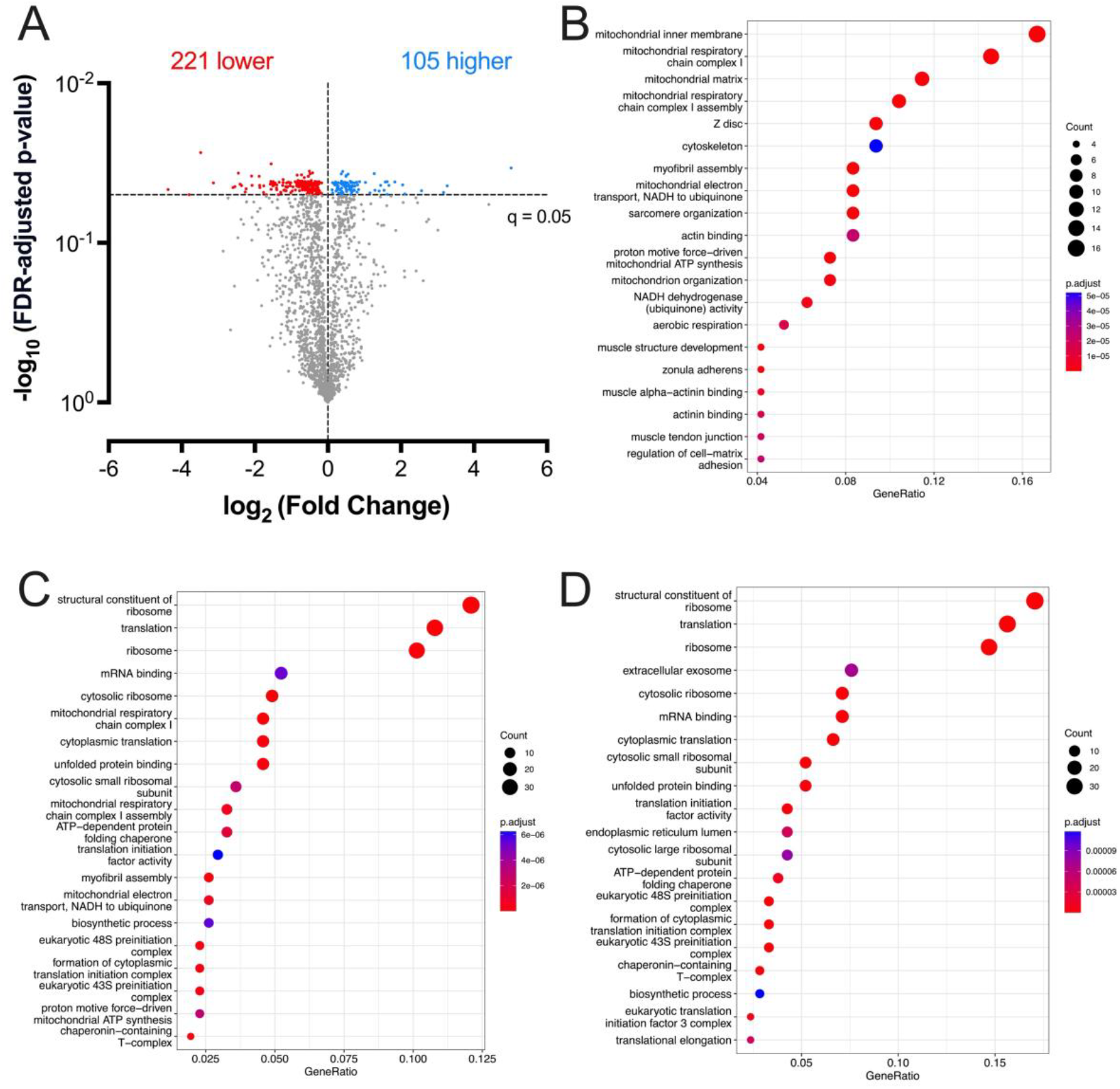
Changes at the protein level following the knockdown of Pc-*prosβ1*. Newly emerging cabbage stem flea beetle (CSFB) adults were fed on oilseed rape leaf disk containing 500 ng of dsProsβ1 or dsmGFP (control dsRNA) and sampled 5 days after the consumption of the leaf disk (n = 5 per group). (a) Volcano plot showing changes in the abundance of proteins in the dsProsβ1 group compared to the dsmGFP group. Permutation-based FDR-adjusted p-value (-log_10_) and fold change (log_2_) are shown for each protein. Blue dots show proteins with significantly higher abundance in dsProsβ1 group and red dots show proteins with significantly lower abundance in the dsProsβ1 group compared to dsmGFP group among the 1980 proteins that were identified and passed filtration. (b-d) The proteins that had significantly higher (b) or lower abundance (c) in the dsProsβ1 group compared to the dsmGFP group were subjected to GO enrichment analysis using R package “ProfileClusterer”. (d) All differentially abundant proteins were also analyzed together.

GO enrichment analysis showed that predominantly mitochondrial and cytoskeleton-related proteins increased in abundance in the dsProsβ group (Fig. 5B). In contrast, the proteins that decreased in abundance in the dsProsβ group, were mostly related to gene expression and translation-associated processes, including ribosome and mRNA binding (Fig. 5C). Interestingly, unfolded protein binding was also enriched among the proteins with lower abundance in the dsProsβ group. Combining all proteins showing differential abundance in the dsProsβ group did not reveal any additional pathways (Fig. 5D).

## 4 DISCUSSION

RNAi has been recognized as a promising alternative to chemical insecticides, and the target gene is one of the most important parameters for effective RNAi-based pest control.^7^ Recently, proteasome has emerged as a very effective RNAi target in multiple insect pests including *L. decemlineata*.^7,11,12^ The commercialization of a proteasome targeting insecticidal dsRNA spray^12^ and the fact that proteasome was one of the top pathways in the genome-wide RNAi-based screen in *Tribolium castaneum*^11^ suggest that dsRNA targeting the proteasome hold great promise in the management of other pests as well. However, a comprehensive understanding of the mechanism of action of proteasome-targeting insecticidal dsRNA was lacking. Here, we combined RISC-bound sRNA-seq, RNA degradome-seq and proteomics to investigate the mechanism of action of dsProsβ1, a proteasome targeting and highly insecticidal dsRNA in CSFB.

The bioinformatics analysis in this study showed that the proteasome subunit *prosβ1* is highly conserved at the protein level among coleopteran pests, which is expected from a gene with such an essential function in the cell. This is also in line with the insight from the genome-wide RNAi screen in *T. castaneum* that almost all effective target genes belong to conserved molecular pathways.^11^ Nonetheless, our previous study showed that certain regions within the effective gene targets transcripts, including *Pc*-*prosβ1*, allow the designing of dsRNA with minimal risk to non-target organisms, including honeybees.^9^

RNA degradome sequencing is yet to be utilized in the context of RNAi-based pest control, which depends on the cleavage of mRNA transcripts in the target pest. We show that a typical pipeline used in plant RNA degradomics^29,31^ can be applied to insect pests to identify major cleavage events caused by insecticidal dsRNA. This has implications for improving RNAi-based pest control because optimizing dsRNA for increased target mRNA cleavages might increase insecticidal efficacy. To that end, we observed that specific nucleotide pairs, such as uracil-uracil (14.3%) and cytosine-adenine (13.8%), occurring at the 10^th^ and 11^th^ positions of the antisense siRNA strand—where they align with the catalytic site of RISC—were associated with increased mRNA cleavage events. This was the case even though most corresponding antisense siRNA reads were relatively low in abundance. Hence, future efforts should focus on an integrated approach where siRNA capable of effectively causing cleavages are also produced from the dsRNA in higher abundance to achieve optimized insecticidal performance. Similar approaches have characterized siRNA sequence features that correlated with high silencing efficiency to be used in human therapeutics.^47–49^ However, it should be noted that accomplishing a similar feat for dsRNA will be more challenging as in the case of dsRNA, we will need to be able to predict both how the dsRNA will be processed into siRNA as well as the efficiencies of each siRNA in the generated siRNA pool.

The discrepancy between the RISC-bound siRNA abundance and the corresponding cleavage events is interesting because the default assumption is that more siRNA would translate into more cleavages at the target site. However, this assumption does not take into account that even if the siRNA is highly abundant, it might not have high efficacy in guiding cleavages due to the biochemical properties of the siRNA.^32^ For instance, the siRNA causing the most observed cleavage event had high GC content at its edges, which might be important for firmly binding to the target mRNA. Subsequently, the sequences at the catalytic site are likely to be important for effectively cleaving the mRNA. It should be noted that the abundance of siRNA processed from dsRNA likely depends on other factors, such as the thermodynamic stability of the siRNA duplex ends, which is important for the guide strand selection process.^32,50,51^ Another potential mechanism leading to this discrepancy between RISC-bound siRNA abundance and cleavage events might be the unloading of siRNA from RISC after effective cleavage events. A similar phenomenon is observed in humans, where the guide RNA bound to the Argonaute-2 is unloaded if the guide RNA is highly complementary to the target RNA.^52^

Interestingly, we observed that more proteins had lower abundance rather than higher abundance upon the inhibition of the proteasome via dsProsβ1. This likely highlights the amino acid recycling function of the proteasome because the limited availability of amino acids causes a reduction in protein synthesis. Supporting this hypothesis, we observed that proteins involved in central dogma, including ribosomal and mRNA binding proteins, had lower abundance in the dsProsβ1group. This is likely the main reason behind the severe mortality caused by the proteasome-targeting dsRNA^9,11,12^ because inhibition of protein synthesis compromises virtually all vital processes in the cell. An interesting enriched process among proteins with lower abundance in the dsProsβ1group was unfolded protein binding. This suggests that inhibition of the proteasome activity induced a negative feedback that reduced the unfolded protein binding activity as well because these processes normally work in tandem.^13^

In contrast, we found that cytoskeleton-related proteins such as actin as well as mitochondrial proteins were increased in abundance upon proteasome inhibition. The turnover of actin filaments is known to be very high because actin plays critical roles in cellular integrity and cellular movements, requiring high plasticity.^53,54^ The enrichment of proteins with high turnover rates is expected upon the inhibition of the proteasome as such proteins are the main targets of proteasome-mediate degradation.^55^ The proteasome is also known to play a crucial role in mitochondrial protein quality control^56,57^, which explains why many mitochondrial proteins increased in abundance upon the inhibition of the proteasome.

## 5 CONCLUSION

In this study, we show that a combination of RNA degradomics, RISC-bound sRNA-seq and proteomics is an insightful approach for investigating the mechanisms of RNAi-based pest control strategies at the molecular level. Particularly, RNA degradomics and RISC-bound sRNA-seq revealed important discrepancies between mature siRNA abundance and corresponding cleavage events. Addressing these discrepancies might help us enhance RNAi-based pest control strategies in the future. Moreover, the proteomics analysis revealed insights into why proteasome-targeting dsRNA causes severe mortality by showing that central dogma-related processes, which are required for all vital functions in the cell, are inhibited, likely due to the reduced amino acid recycling.

## ACKNOWLEDGEMENTS

Doga Cedden and Gözde Güney were funded by the Deutscher Akademischer Austauschdienst research grants program during the project. Open Access funding enabled and organized by Projekt DEAL. The authors thank Kerstin Schmitt, Oliver Valerius, Johan Zicola, and Gregor Bucher for their help in the study. Some of the figures were created with BioRender.com.

## AUTHOR CONTRIBUTIONS

DC conceived the ideas. DC, GG, SS, and MR and designed the methodology. DC and GG performed the experiments and analyzed the data. DC drafted the manuscript. DC, GG, MR and SS edited and revised the manuscript.

## DATA AVAILABILITY STATEMENT

The RISC-bound sRNA-seq data (BioProject: PRJNA1157760) and the RNA degradome sequencing data (PRJNA1196165) were deposited into the NCBI Sequence Read Archive. Processed data sets are provided in the Supporting Information 2. The protein quantification data was made available publicly on FigShare (https://doi.org/10.6084/m9.figshare.27939765.v1). The mass spectrometry proteomics data have been deposited to the ProteomeXchange Consortium via the PRIDE partner repository with the dataset identifier PXD061878. Further details can be obtained from the corresponding author.

## References

1 Deutsch CA, Tewksbury JJ, Tigchelaar M, Battisti DS, Merrill SC, Huey RB, et al., Increase in crop losses to insect pests in a warming climate, Science 361:916–919, American Association for the Advancement of Science (2018).

2 Feyereisen R, Molecular biology of insecticide resistance, Toxicology Letters 82– 83:83–90 (1995).

3 Sparks TC and Nauen R, IRAC: Mode of action classification and insecticide resistance management, Pestic Biochem Physiol 121:122–128 (2015).

4 Sparks TC, Crossthwaite AJ, Nauen R, Banba S, Cordova D, Earley F, et al., Insecticides, biologics and nematicides: Updates to IRAC’s mode of action classification - a tool for resistance management, Pesticide Biochemistry and Physiology 167:104587 (2020).

5 Chen Y and De Schutter K, Biosafety aspects of RNAi-based pests control, Pest Management Science 80:3697–3706 (2024).

6 Baum JA, Bogaert T, Clinton W, Heck GR, Feldmann P, Ilagan O, et al., Control of coleopteran insect pests through RNA interference, Nat Biotechnol 25:1322–1326, Nature Publishing Group (2007).

7 Cedden D and Bucher G, The quest for the best target genes for RNAi-mediated pest control, Insect Molecular Biology n/a (2024).

8 Mehlhorn S, Hunnekuhl VS, Geibel S, Nauen R, and Bucher G, Establishing RNAi for basic research and pest control and identification of the most efficient target genes for pest control: a brief guide, Front Zool 18:60 (2021).

9 Cedden D, Güney G, Debaisieux X, Scholten S, Rostás M, and Bucher G, Effective target genes for RNA interference-based management of the cabbage stem flea beetle, Insect Molecular Biology n/a (2024).

10 Knorr E, Fishilevich E, Tenbusch L, Frey MLF, Rangasamy M, Billion A, et al., Gene silencing in Tribolium castaneum as a tool for the targeted identification of candidate RNAi targets in crop pests, Sci Rep 8:2061, Nature Publishing Group (2018).

11 Buer B, Dönitz J, Milner M, Mehlhorn S, Hinners C, Siemanowski-Hrach J, et al., Superior target genes and pathways for RNAi-mediated pest control revealed by genome-wide analysis in the beetle, Pest Management Science 81:1026–1036 (2025).

12 Rodrigues TB, Mishra SK, Sridharan K, Barnes ER, Alyokhin A, Tuttle R, et al., First Sprayable Double-Stranded RNA-Based Biopesticide Product Targets Proteasome Subunit Beta Type-5 in Colorado Potato Beetle (Leptinotarsa decemlineata), Frontiers in Plant Science 12 (2021).

13 Marshall RS and Vierstra RD, Dynamic Regulation of the 26S Proteasome: From Synthesis to Degradation, Frontiers in Molecular Biosciences 6 (2019).

14 Tanaka K, The proteasome: Overview of structure and functions, Proc Jpn Acad Ser B Phys Biol Sci 85:12–36 (2009).

15 Obbard DJ, Gordon KHJ, Buck AH, and Jiggins FM, The evolution of RNAi as a defence against viruses and transposable elements, Philos Trans R Soc Lond B Biol Sci 364:99–115 (2009).

16 Galiana-Arnoux D, Dostert C, Schneemann A, Hoffmann JA, and Imler J-L, Essential function in vivo for Dicer-2 in host defense against RNA viruses in drosophila, Nat Immunol 7:590–597 (2006).

17 Wang X-.H, RNA interference directs innate immunity against viruses in adult Drosophila, Science 312 (2006).

18 Lim DH, Kim J, Kim S, Carthew RW, and Lee YS, Functional analysis of dicer-2 missense mutations in the siRNA pathway of Drosophila, Biochem Biophys Res Commun 371 (2008).

19 Tijsterman M and Plasterk RHA, Dicers at RISC: The Mechanism of RNAi, Cell 117:1–3 (2004).

20 Zhu KY and Palli SR, Mechanisms, Applications, and Challenges of Insect RNA Interference, Annual review of entomology 65:293 (2019).

21 Iwakawa H-. O and Tomari Y, Life of RISC: Formation, action, and degradation of RNA-induced silencing complex, Mol Cell 82 (2021).

22 Kawamata T and Tomari Y, Making RISC, Trends Biochem Sci 35:368–376 (2010).

23 Khvorova A, Reynolds A, and Jayasena SD, Functional siRNAs and miRNAs exhibit strand bias, Cell 115 (2003).

24 Warner DJ, Allen-Williams LJ, Warrington S, Ferguson AW, and Williams IH, Mapping, characterisation, and comparison of the spatio-temporal distributions of cabbage stem flea beetle (Psylliodes chrysocephala), carabids, and Collembola in a crop of winter oilseed rape (Brassica napus), Entomologia Experimentalis et Applicata 109:225–234 (2003).

25 Li Z, Costamagna AC, Beran F, and You M, Biology, Ecology, and Management of Flea Beetles in Brassica Crops, Annual Review of Entomology 69:199–217 (2024).

26 Ortega-Ramos PA, Coston DJ, Seimandi-Corda G, Mauchline AL, and Cook SM, Integrated pest management strategies for cabbage stem flea beetle (Psylliodes chrysocephala) in oilseed rape, GCB Bioenergy 14:267–286 (2022).

27 Højland DH, Nauen R, Foster SP, Williamson MS, and Kristensen M, Incidence, Spread and Mechanisms of Pyrethroid Resistance in European Populations of the Cabbage Stem Flea Beetle, Psylliodes chrysocephala L. (Coleoptera: Chrysomelidae), PLOS ONE 10:e0146045, Public Library of Science (2015).

28 Cedden D, Güney G, Scholten S, and Rostás M, Lethal and sublethal effects of orally delivered double-stranded RNA on the cabbage stem flea beetle, Psylliodes chrysocephala, Pest Management Science 80:2282–2293 (2024).

29 Ma X, Yin X, Tang Z, Ito H, Shao C, Meng Y, et al., The RNA degradome: a precious resource for deciphering RNA processing and regulation codes in plants, RNA Biology 17:1223 (2020).

30 Jackowiak P, Nowacka M, Strozycki PM, and Figlerowicz M, RNA degradome—its biogenesis and functions, Nucleic Acids Research 39:7361–7370 (2011).

31 Li Y-F, Zhao M, Wang M, Guo J, Wang L, Ji J, et al., An improved method of constructing degradome library suitable for sequencing using Illumina platform, Plant Methods 15:134 (2019).

32 Cedden D, Guney G, Rostas M, and Bucher G, Optimizing dsRNA sequences for RNAi in pest control and research with the dsRIP Web-Platform, Cold Spring Harbor Laboratory (2025).

33 Güney G, Cedden D, Körnig J, Ulber B, Beran F, Scholten S, et al., Physiological and transcriptional changes associated with obligate aestivation in the cabbage stem flea beetle (Psylliodes chrysocephala), Insect Biochemistry and Molecular Biology 173:104165 (2024).

34 Jumper J, Highly accurate protein structure prediction with AlphaFold, Nature 596 (2021).

35 Abramson J, Adler J, Dunger J, Evans R, Green T, Pritzel A, et al., Accurate structure prediction of biomolecular interactions with AlphaFold 3, Nature 630:493–500, Nature Publishing Group (2024).

36 Wang J, Chitsaz F, Derbyshire MK, Gonzales NR, Gwadz M, Lu S, et al., The conserved domain database in 2023, Nucleic Acids Res 51:D384–D388 (2023).

37 Tamura K, Stecher G, and Kumar S, MEGA11: Molecular Evolutionary Genetics Analysis Version 11, Molecular Biology and Evolution 38:3022–3027 (2021).

38 Langmead B, Trapnell C, Pop M, and Salzberg SL, Ultrafast and memory-efficient alignment of short DNA sequences to the human genome, Genome Biology 10:R25 (2009).

39 Untergasser A, Cutcutache I, Koressaar T, Ye J, Faircloth BC, Remm M, et al., Primer3—new capabilities and interfaces, Nucleic Acids Research 40:e115 (2012).

40 Güney G, Schmitt K, Zicola J, Toprak U, Rostás M, Scholten S, et al., The MicroRNA pathway regulates obligatory aestivation in a flea beetle, Cold Spring Harbor Laboratory (2025).

41 Grentzinger T, Oberlin S, Schott G, Handler D, Svozil J, Barragan-Borrero V, et al., A universal method for the rapid isolation of all known classes of functional silencing small RNAs, Nucleic Acids Res 48:e79 (2020).

42 Rappsilber J, Mann M, and Ishihama Y, Protocol for micro-purification, enrichment, pre-fractionation and storage of peptides for proteomics using StageTips, Nature Protocols 2:1896–1906, Nature Publishing Group (2007).

43 J R, Y I, and M M, Stop and go extraction tips for matrix-assisted laser desorption/ionization, nanoelectrospray, and LC/MS sample pretreatment in proteomics, PubMed (2003).

44 Tyanova S, Temu T, Sinitcyn P, Carlson A, Hein MY, Geiger T, et al., The Perseus computational platform for comprehensive analysis of (prote)omics data, Nature Methods 13:731–740, Nature Publishing Group (2016).

45 Murari E, Meadows D, Cuda N, and Mangone M, A comprehensive analysis of 3′UTRs in Caenorhabditis elegans, Nucleic Acids Research 52:7523–7538 (2024).

46 Yang W, Hsu PL, Yang F, Song J-E, and Varani G, Reconstitution of the CstF complex unveils a regulatory role for CstF-50 in recognition of 3′-end processing signals, Nucleic Acids Research 46:493–503 (2018).

47 Han Y, He F, Chen Y, Liu Y, and Yu H, SiRNA silencing efficacy prediction based on a deep architecture, BMC Genomics 19:669 (2018).

48 Reynolds A, Leake D, Boese Q, Scaringe S, Marshall WS, and Khvorova A, Rational siRNA design for RNA interference, Nat Biotechnol 22:326–330, Nature Publishing Group (2004).

49 Kurreck J, siRNA Efficiency: Structure or Sequence—That Is the Question, J Biomed Biotechnol 2006:83757 (2006).

50 Malefyt AP, Wu M, Vocelle DB, Kappes SJ, Lindeman SD, Chan C, et al., Improved asymmetry prediction for siRNAs, FEBS J 281:320–330 (2014).

51 Schwarz DS, Asymmetry in the assembly of the RNAi enzyme complex, Cell 115 (2003).

52 De N, Young L, Lau P-W, Meisner N-C, Morrissey DV, and MacRae IJ, Highly Complementary Target RNAs Promote Release of Guide RNAs from Human Argonaute2, Molecular cell 50:344 (2013).

53 Onishi M, Pecani K, Jones T, Pringle JR, and Cross FR, F-actin homeostasis through transcriptional regulation and proteasome-mediated proteolysis, Proceedings of the National Academy of Sciences 115:E6487–E6496, Proceedings of the National Academy of Sciences (2018).

54 Goode BL, Eskin J, and Shekhar S, Mechanisms of actin disassembly and turnover, The Journal of Cell Biology 222:e202309021 (2023).

55 Bartolome A, Heiby JC, Di Fraia D, Heinze I, Knaudt H, Spaeth E, et al., Quantitative mapping of proteasome interactomes and substrates using ProteasomeID, ed. by Lehrbach N and Cooper JA, eLife 13:RP93256, eLife Sciences Publications, Ltd (2024).

56 Rödl S and Herrmann JM, The role of the proteasome in mitochondrial protein quality control, IUBMB Life 75:868–879 (2023).

57 Heo J-M and Rutter J, Ubiquitin-dependent mitochondrial protein degradation, The international journal of biochemistry & cell biology 43:1422 (2011).

