## Supporting information 1 for "RNA degradomics and proteomics reveal the mechanism of dsProsβ1-mediated proteasome targeting in the cabbage stem flea beetle"

**Supporting Table 1. The nucleic acid and translated amino acid sequences of *Pc*-*prosβ1*. The region targeted by dsProsβ1 is underlined.**

>Nucleic acid sequence of *Pc*-*prosβ1*, TRINITY_DN20894_c0_g1

TTTCGGCTTGTCATCTTTATAATATCTTGTCATTCGATATTCAATATCATTGTGTTTTTATTTTCCATTTTATTCATACGTATTGTTACTAAAGAGTTGATATTATAAACCTTAATTTTATCATTCATGATGATGCAAACAGAAATGAATGCTTATTATCCTGACTATTCTACAGCTCCAATCCAACGACAAATTAACCCCTATGCTGATAATGGAGGGAGTGTTGTAGCGATAGCAGGAAATGATTTCGTAGTTGTGGGAGCTGATACTCGTCTCAGTTCAGGTTTTTCAATTTACACAAGAAACCAAAATAAATTATTTCCTATGTCCGATACAACTGTATTAGGTTGTGCAGGATGTTGGTGCGACACTTTAACTTTAACCCGAATTTTGAAATCTCGTATGCAGATGTACCAGCAGGAACATAACAAACAAATGTCTACAACAGCATGTGCACAGATGTTATCCACCATGCTGTATTATAAGAGATTTTTCCCTTACTATATATCCAATATACTTGTTGGATTGGATAATGAAGGAAAAGGTTGTGTTTATAGCTATGATCCAATTGGGCATTGTGAAAAGGCCACCTACAGAGCAGGTGGCTCTGCAGGTGCATTACTTCAACCTCTTTTGGACAACCAGATTGGACAAAAAAATCTGGTGAAACCATCATCTGAAGATCTTTCTCAAGAGAAAGCTCTTGCTATATTGAAAGATGTATTCATTGCTGCTGCAGAGAGAGACATCTATACTGGAGATAGTGTTCTTATCAATATTATTACAAAGGATGGAACTAAAGTAGAAACTTTTGAATTAAGAAAAGATTAATATTTTGTATTATTTTTTAAATAAAGCATTCAATTATTCATTTCTTTCAAAGGTCATTTGATTTTTTATTTATTAAGAAGATCCCTTGCTGGATATTAAAAAGGTTGTAGATTATATTACATAGCCAAAAAAAAAACCTGTAAAATTTAAGGAAACCAATCAAAATCGTGAGTATATTGTTTATTAAAAAAACCTGTAAAATTTAAGGAAACCAATCAAAATCGTGAGTATATTGTTTATTAAAAAAA

>Translated amino acid sequence of Pc-Prosβ1

MMMQTEMNAYYPDYSTAPIQRQINPYADNGGSVVAIAGNDFVVVGADTRLSSGFSIYTRNQNKLFPMSDTTVLGCAGCWCDTLTLTRILKSRMQMYQQEHNKQMSTTACAQMLSTMLYYKRFFPYYISNILVGLDNEGKGCVYSYDPIGHCEKATYRAGGSAGALLQPLLDNQIGQKNLVKPSSEDLSQEKALAILKDVFIAAAERDIYTGDSVLINIITKDGTKVETFELRKD

^1^Accession for CSFB adult transcriptome is GKIH00000000.1

**Supporting Table 2. Sequences of the primers and adaptors used in the study**

| Name | Purpose | Direction | Sequence |
| --- | --- | --- | --- |
| dsmGFP | In vitro transcription | Forward with T7 | GAATTGTAATACGACTCACTATAGGACCCTGACCTACGGCCTAT |
| dsmGFP | In vitro transcription | Reverse  with T7 | GAATTGTAATACGACTCACTATAGGTGCCGTCCTCGTACTAGTT |
| dsProsβ1 | In vitro transcription | Forward with T7 | GAATTGTAATACGACTCACTATAGGAGGATGTTGGTGCGACACTT |
| dsProsβ1 | In vitro transcription | Reverse  with T7 | GAATTGTAATACGACTCACTATAGGATGTCTCTCTCTGCAGCAGC |
| dsmGFP | In vitro transcription | Forward | GGACCCTGACCTACGGCCTAT |
| dsmGFP | In vitro transcription | Reverse | GGTGCCGTCCTCGTACTAGTT |
| dsProsβ1 | In vitro transcription | Forward | AGGATGTTGGTGCGACACTT |
| dsProsβ1 | In vitro transcription | Reverse | ATGTCTCTCTCTGCAGCAGC |
| 5′ RNA adaptor | RNA degradome | Forward | GUUCAGAGUUCUACAGUCCGACGAUCAGCAG |
| RT-primer | RNA degradome | Reverse | CGAGCACAGAATTAATACGACTTTTTTTTTTTTTTTTTT |
| 5′ adaptor | RNA degradome | Forward | GTTCAGAGTTCTACAGTCCGAC |
| 3′ adaptor | RNA degradome | Reverse | CGAGCACAGAATTAATACGACT |
| dsDNA top | RNA degradome | Forward | NNTGGAATTCTCGGGTGCCAAGG |
| dsDNA bottom | RNA degradome | Reverse | CCTTGGCACCCGAGAATTCCA |
| Final 5′PCR primer | RNA degradome | Forward | AATGATACGGCGACCACCGAGATCTACACGTTCAGAGTTCTACAGTCCGA |
| Final 3′PCR primer for dsProsβ1-fed CSFB | RNA degradome | Reverse with index | CAAGCAGAAGACGGCATACGAGATACATCGGTGACTGGAGTTCAGACGTGTGCTCTTCCGATCT* |
| Final 3′PCR primer for mGFP-fed CSFB | RNA degradome | Reverse with index | CAAGCAGAAGACGGCATACGAGATCGTGATGTGACTGGAGTTCAGACGTGTGCTCTTCCGATCT* |

* The index sequence is underlined.

**Supporting Table 3. miRanda (www.microrna.org) target prediction results for a miRNA potentially targeting the 3’ UTR region of *Pc-prosβ1***

| miRNA | Target gene | miRanda score | Energy-Kcal/Mol | Aligned region | Alignment length | Mature sequence |
| --- | --- | --- | --- | --- | --- | --- |
| OV651826.1_41984 | TRINITY_DN20894_c0_g1  *Pc-prosβ1* | 157 | -10.25 | 142-165 from 3’ UTR start | 19 | UUUUGAUUGUUGCUCAGAAAGC |

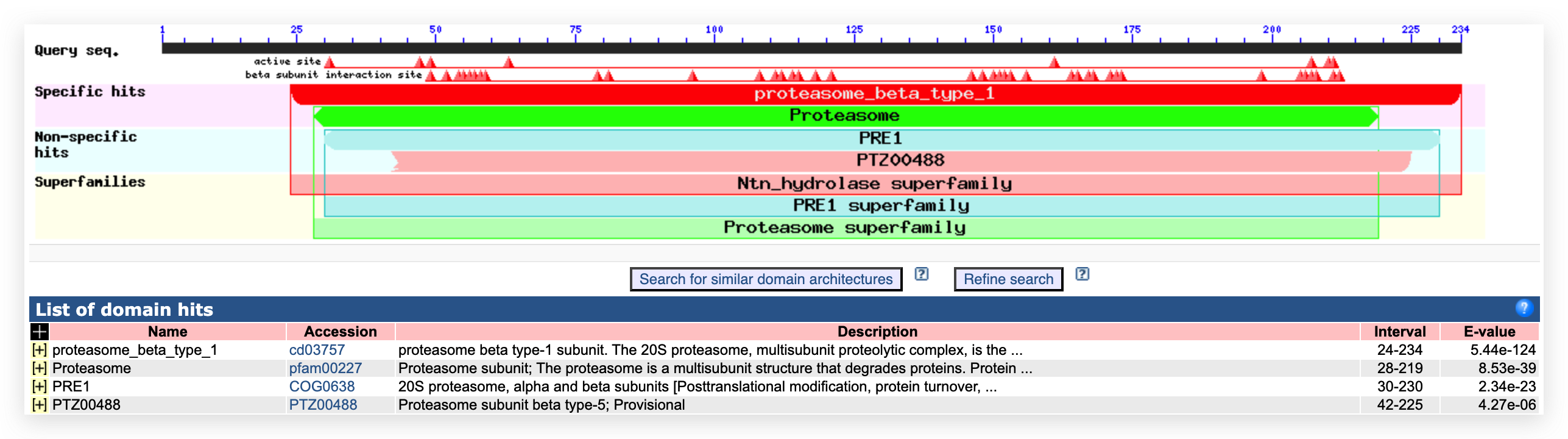

**Supporting Figure 1. Conserved domain analysis of Pc-Prosβ1.** Pc-prosβ1 contains an ORF spanning 702 nucleotides and encodes for 234 amino acids.Conserved domain analysis suggested N-terminal nucleophile hydrolase (Ntn hydrolase) activity for Pc-Prosβ1, which is expected of a Beta 1 subunit.
